# Cancer-Associated Fibroblasts Promote Prostate Cancer Malignancy via Metabolic Rewiring and Mitochondrial Transfer

**DOI:** 10.1101/558080

**Authors:** Luigi Ippolito, Andrea Morandi, Maria Letizia Taddei, Matteo Parri, Giuseppina Comito, Alessandra Iscaro, Maria Rosaria Raspollini, Francesca Magherini, Elena Rapizzi, Julien Masquelier, Giulio G. Muccioli, Pierre Sonveaux, Paola Chiarugi, Elisa Giannoni

**Affiliations:** Department of Experimental and Clinical Biomedical Sciences, University of Florence, 50134 Florence, Italy; Department of Experimental and Clinical Medicine, University of Florence, 50134 Florence, Italy; Histopathology and Molecular Diagnostics, University Hospital Careggi, Largo Brambilla, 3, 50134 Florence, Italy; Bioanalysis and Pharmacology of Bioactive Lipids Research Group, Louvain Drug Research Institute (LDRI), Université catholique de Louvain (UCL), B-1200 Brussels, Belgium; Pole of Pharmacology, Institut de Recherche Expérimentale et Clinique (IREC), Université catholique de Louvain (UCL), B-1200 Brussels, Belgium; Tuscany Tumour Institute (ITT) and Excellence Centre for Research, Transfer and High Education DenoTHE, Florence, Italy.

**Keywords:** OXPHOS, sirtuins, cancer-associated fibroblasts (CAFs), mitochondrial transfer, prostate cancer

## Abstract

Cancer-associated fibroblasts (CAFs) are the major cellular stromal component of many solid tumors. In prostate cancer (PCa), CAFs establish a metabolic symbiosis with PCa cells, contributing to cancer aggressiveness through lactate shuttle. In this study, we report that lactate uptake alters the NAD^+^/NADH ratio in the cancer cells, which culminates with SIRT1-dependent PGC-1α activation and subsequent enhancement of mitochondrial mass and activity. The high exploitation of mitochondria results in tricarboxylic acid cycle deregulation, accumulation of oncometabolites and in the altered expression of mitochondrial complexes, responsible for superoxide generation. Additionally, cancer cells hijack CAF-derived functional mitochondria through the formation of cellular bridges, a phenomenon that we observed in both *in vitro* and *in vivo* PCa models. Our work reveals a crucial function of tumor mitochondria as the energy sensors and transducers of CAF-dependent metabolic reprogramming and underscores the reliance of PCa cells on CAF catabolic activity and mitochondria trading.

## INTRODUCTION

Fibroblasts are the major constituent of the stromal compartment of many solid cancers, and their activation to a cancer-associated fibroblast (CAF) phenotype contributes to tumor initiation and progression. We and others have showed that CAFs sustain tumor progression by reprogramming cancer cell metabolism [1, 2]. In PCa, both CAFs and tumor cells undergo a reciprocal metabolic reprogramming and establish a symbiotic relationship. CAFs are converted to aerobic glycolysis when in contact with cancer cells, that reprogramme to oxidative phosphorylation (OXPHOS) upon CAF-released lactate upload via the monocarboxylate transporter 1 (MCT1). Lactate-driven conversion of PCa cells to OXPHOS is sustained by the nuclear translocation of pyruvate kinase isoform M2 (PKM2), where it acts as a transcriptional regulator [3]. CAF-induced dependence on mitochondrial respiration of cancer cells further confers chemotherapy resistance [4], promotes redox-dependent epithelial-to-mesenchymal transition (EMT) and increases metastatic burden [3].

It is now emerging that mitochondria are essential in cancer initiation and progression [5] and consequently OXPHOS has been associated with metastatic potential and resistance to therapy in many cancers [4, 6, 7]. Additionally, important mitochondrial metabolism regulators like the transcriptional coactivator peroxisome proliferator-activated receptor-gamma coactivator-1 (PGC1-α) have been reported to support metastatic dissemination in melanoma and breast cancer [8]. However, mitochondrial metabolism has been also described to be detrimental in a model of prostate tumor progression [9], highlighting that microenvironmental cues and stromal cells within the tumor microenvironment can be crucial to better understand the role of this fascinating organelle. In this context, the transfer of ‘intact’ functional mitochondria and/or mitochondrial DNA (mtDNA) from stromal to cancer cells has been reported to proficiently restore OXPHOS in mtDNA-deprived cancer cells, hence re-establishing tumor-initiating efficacy [10–13].

Here, we investigate how CAFs regulate mitochondrial dynamics in PCa cells. Crucially, we demonstrate that molecular and metabolic inputs generated by CAFs are sensed by the SIRT1/PGC-1α axis of the cancer cells that lead to mitochondria reshape. Additionally, we provide evidence that CAFs donate their dispensable but functional mitochondria to the cancer cells leading to an enhancement of their malignant phenotype. Understanding the role of mitochondria in this PCa model of metabolic symbiosis offers an opportunity to eventually target tumor:stroma co-evolution and co-operation.

## RESULTS

### CAF contact activates SIRT1/PGC-1α and potentiates mitochondrial metabolism in PCa cells

To investigate the molecular players involved in the metabolic conversion of PCa cells towards OXPHOS, we initially focused on sirtuins (SIRTs), NAD^+^-dependent deacetylases that act as sensors of nutrient availability, whose expression being increased upon nutrient deprivation [14, 15]. Indeed, after 24 hours of PC3 cells exposure to CAF-conditioned medium (CM), we observed a significant increase in NADH levels (**Fig. 1A, left**) compared to HPF-CM-treated cells, possibly due to the oxidation of the lactate present in the CAF-CM. After 48 and 72 hours NADH levels return to basal, whereas NAD^+^ levels increase (**Fig. 1A, right and Fig. S1A**), as a consequence of electron transport chain (ETC) enhanced activity. Notably, the NAD^+^ and NADH levels were unaffected when cancer cells were exposed to CM from healthy human prostate fibroblasts (HPFs). SIRTs expression has been previously reported to be dependent on the NAD^+^/NADH ratio [16]. Accordingly, western blot analysis revealed a rapid NADH-dependent increase in SIRT1 expression levels (**Fig. 1B**) and subsequent NAD^+^-dependent SIRT1 activation at 48 and 72 hours after CAF-conditioning, as revealed by the deacetylation of PGC-1α (**Fig. 1B**), a regulator of mitochondrial biogenesis and OXPHOS [17] that is an established target of SIRT1 [18]. The link between SIRT1 activity and PGC-1α deacetylation (*i.e.* activation) is supported by the negative correlation between NAD^+^ levels and PGC-1α acetylation status (r_s_=-0.49; *P*<0.0001) (**Fig. 1C**) and further strengthened by the impairment of CAF-induced PGC-1α deacetylation upon SIRT1 silencing (**Fig. S1B**). The relevant role of lactate in mediating the early rise of NADH levels (24 hours) upon CAF-CM administration was reinforced by the inhibitory effect exerted by the inward lactate transporter MCT1 inhibitor (AR-C155858) and by the ability of exogenous lactate administration to resemble CAF conditioning (**Fig. 1D**). We also observed that lactate was the only metabolite that was able to induce a significant SIRT1 overexpression (**Fig. 1E**) and that lactate-induced SIRT1 upregulation and PGC-1α deacetylation were impaired by preventing lactate upload by cancer cells **(Fig. 1F)**, highlighting an additional role of lactate beyond its function as a nutrient.

**Figure 1.**
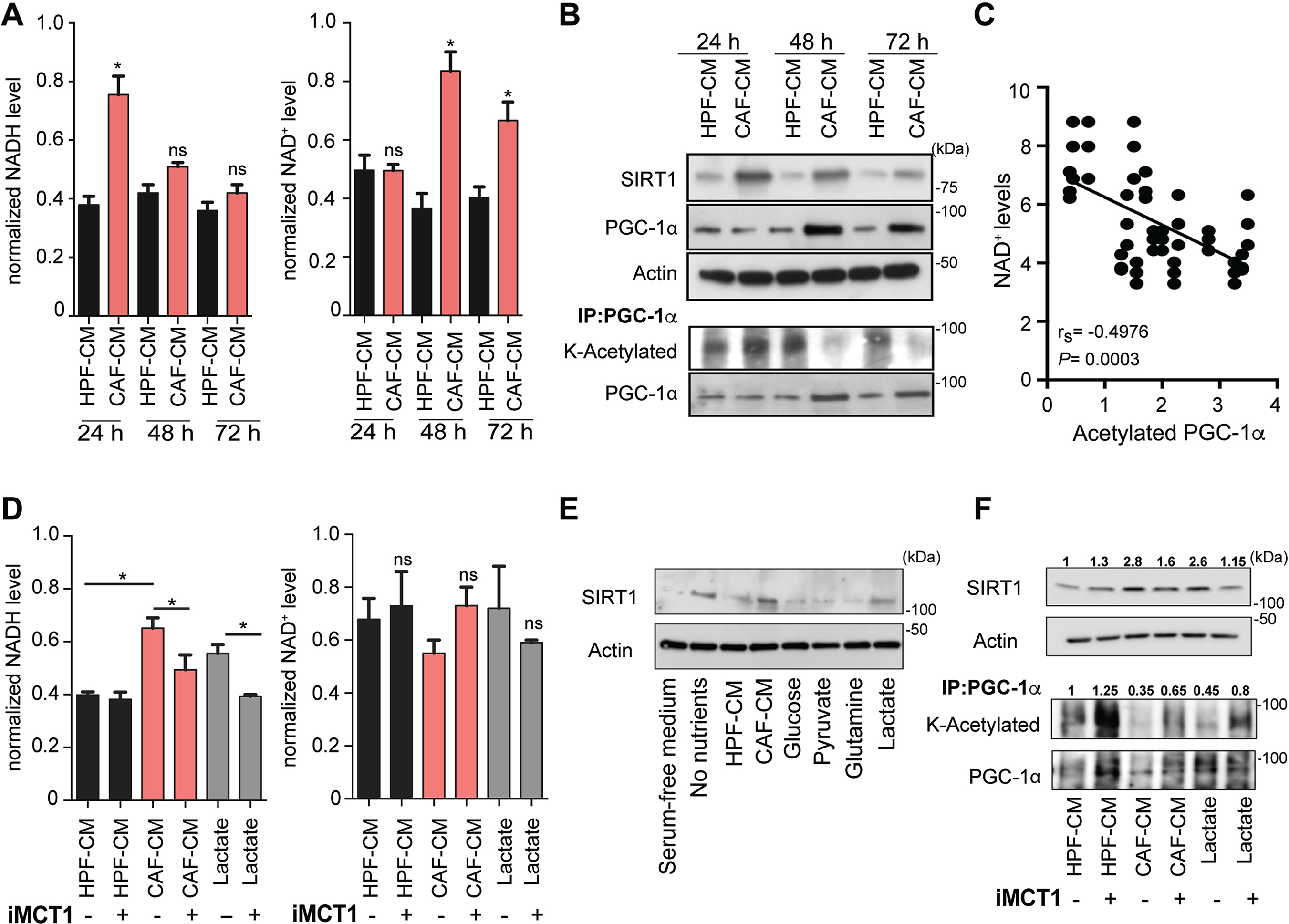
CAFs conditioning triggers a lactate-dependent SIRT1/PGC-1α axis activation in PCa cells. PC3 cells were cultured with conditioned medium (CM) from HPF or CAF for 24, 48 or 72 hours. **A.** Time-point of NADH (left plot) and NAD^+^ (right plot) values, normalized on protein content, have been plotted. Data represent mean ± SEM, *n* = 3. ns, no significant *P < 0.05, by two-tailed *t* test. **B.** Total cell lysates or PGC-1α immunoprecipitates (IP) were subjected to Western blot (WB) analysis as indicated. **C**. Pearson’s correlation scatter plot of the NAD^+^ levels and acetylation status of PGC-1α in PC3 cells treated with CAF-CM for 48 h. **D.** NADH and NAD^+^ normalized levels. PC3 cells were exposed to HPF-CM, CAF-CM or 10mM lactate for 24h, with or without MCT1 inhibitor (AR-C155858, 10µM) administration to. **E**. PC3 cells were treated for 16 h in glucose/pyruvate/glutamine-free medium (‘no nutrients’, used as positive control) and 10 mM lactate, 2 mM glutamine, 1 mM pyruvate and 25 mM glucose where then administered for 24 h. SIRT1 expression was assessed by WB. **F**. SIRT1 expression and PGC-1α acetylation were assessed on PC3 cell lysates and PGC-1α IP, respectively. Quantification analysis of acetylation and SIRT1 expression was relative to their respective controls (the fold change numbers were indicated).

The CAF-mediated activation of the SIRT1/PGC-1α axis has a clear impact on mitochondrial metabolism, has revealed by the increase in mitochondrial activity in both CAF-CM exposed PC3 and DU145 cells, as revealed by FACS and confocal analyses (**Fig. 2 A-B**). Notably, CAF conditioning induce a significant increase in mitochondrial mass (**Fig. 2C**), which likely contributes to the improvement of the overall mitochondrial metabolism. These observation were corroborated by an increase in oxygen consumption rate (OCR) (**Fig. 2D**), confirming the role of CAFs in promoting PCa cells mitochondrial respiration. To investigate the impact of CAF-mediated mitochondrial reprogramming on the tricarboxylic acid (TCA) cycle, PC3 cells exposed to CAF-CM or to HPF-CM were subjected to liquid chromatography mass spectrometry (LC-MS) (**Fig. 2E**). Comparative analysis revealed a significant increase of citrate, succinate, fumarate and malate content in CAF-conditioned samples (**Fig. 2E**), whereas α-ketoglutarate (α-KG) and 2-hydroxyglutarate levels were unaffected by CAF exposure.

**Figure 2.**
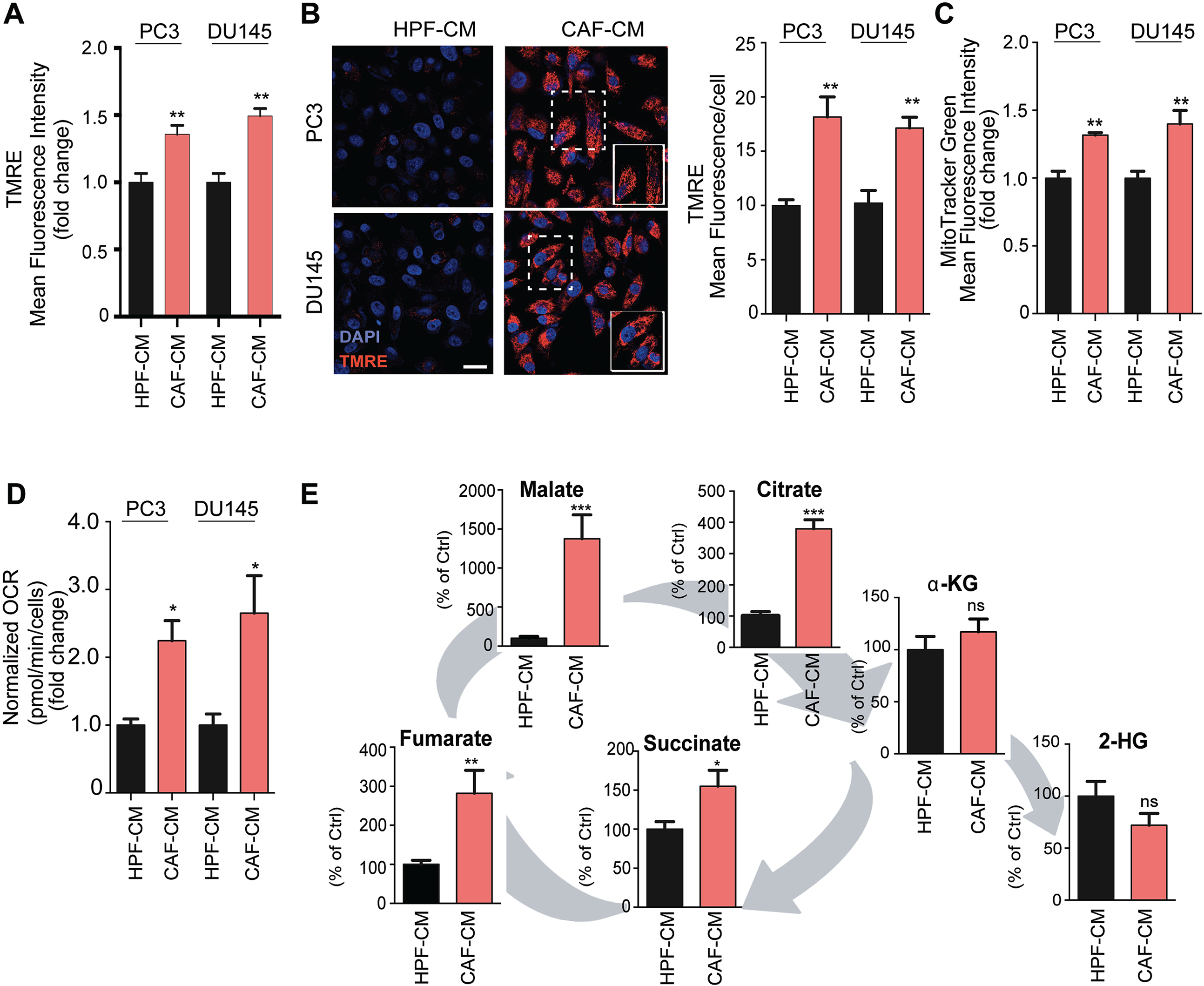
CAFs promote a functional rewiring of mitochondria in PCa cells. **A-C**. Mitochondrial membrane potential (TMRE) and mitochondrial mass (MitoTracker Green) of PCa cells were measured by FACS analysis after 48 hours treatment with HPF-or CAF-CM. Confocal images of TMRE/DAPI-stained cells were also shown, with the corresponding quantification plot. Scale bar, 5 µm (E). **D**. Measurement of oxygen consumption rate (OCR, pmol O_2_/min/cells) in PCa cells, exposed to HPF-CM or CAF-CM for 48 hours. Data represent mean ± SEM, *n =* 3. ns, no significant, *P < 0.05; **P < 0.005; ***P < 0.001, by two-tailed *t* test. **E**. LC-MS analysis of the TCA cycle metabolites. Data were represented as percentage compared to the control. *n* = 3. ns, no significant, *P < 0.05; **P < 0.005; ***P < 0.001, by two-tailed *t* test.

### CAF-induced SIRT1/PGC-1α activation drives PCa cell invasion

Mitochondrial biogenesis and PGC-1α-dependent OXPHOS metabolism are implicated in breast cancer cell motility and metastasis [8, 10]. Therefore, we investigated whether CAF-induced mitochondrial enhancement is associated with PCa cells invasiveness. SIRT1 silencing impaired the ability of PC3 cells to undergo EMT (**Fig. 3A**) and to invade (**Fig. 3B**) following CAF conditioning. Additionally, SIRT1 silencing impaired CAF-induced hypoxia inducible factor-1α (HIF-1α) stabilization in PC3 and DU145 cells (**Fig. 3A**), a well-known CAF-induced malignant feature of PCa cells [3, 19]. Similar results were obtained in DU145 cells (**Fig. S1C**). CAF-induced EMT and invasive abilities of PC3 cells (**Fig. 3C-D**), as well as their mitochondrial improvement (**Fig. 3E**), were also negatively affected by PGC-1α silencing. The importance of the SIRT1/PGC-1α pathway in sustaining the CAF-induced mitochondrial function improvement was strengthened by the negative impact of SIRT1 and PGC-1α silencing on the oxygen consumption rate (**Fig. 3F** and **Fig. S1D**). The impairment of SIRT1 upregulation also prevented TCA cycle rearrangement and the accumulation of citrate, succinate, fumarate and malate upon CAF-CM exposure (**Fig. 3G**).

**Figure 3.**
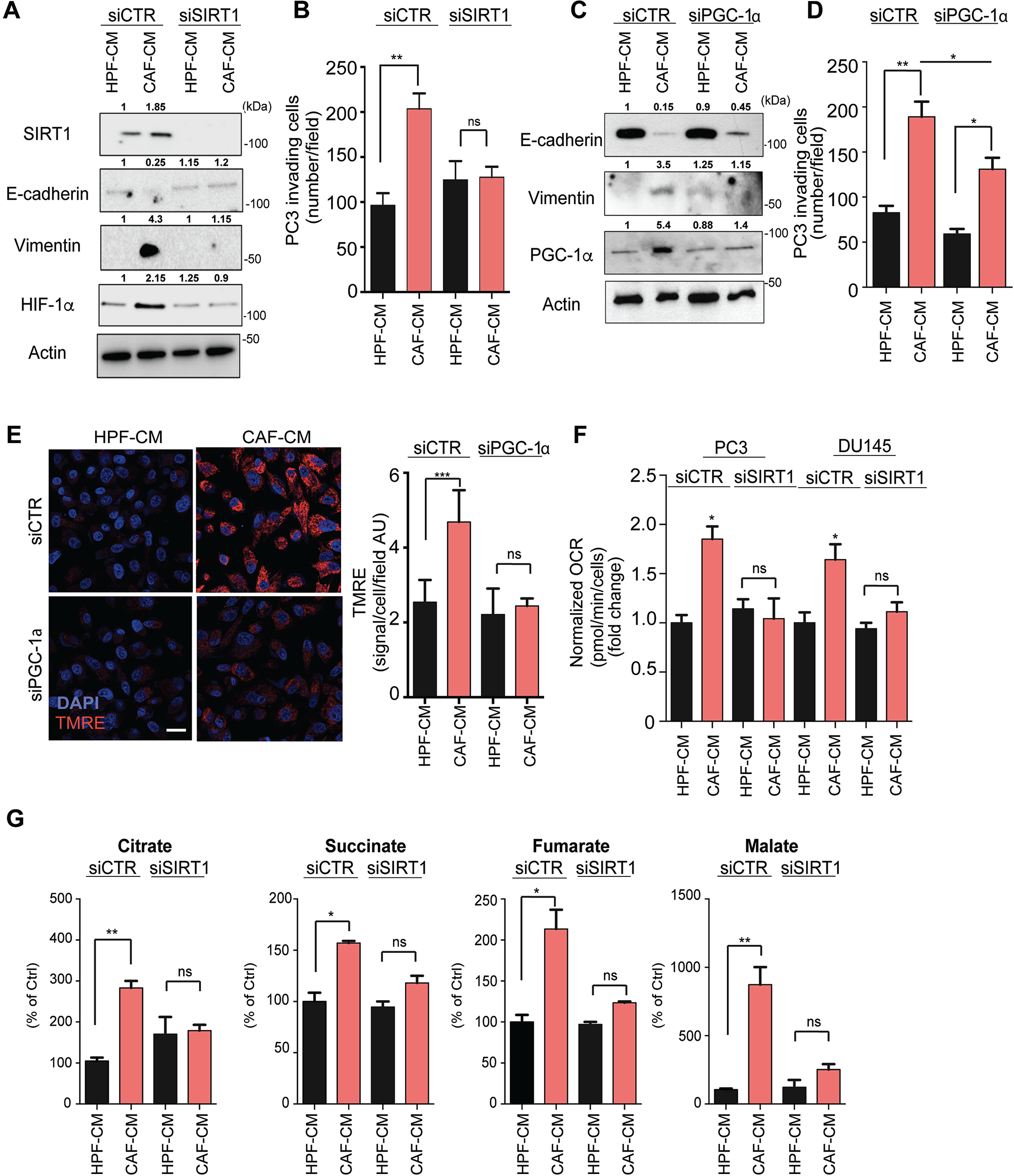
CAFs-mediated SIRT1/PGC-1α axis activation is crucial for invasive traits of PCa cells. PC3 or DU145 cells were silenced with non-targeting control (siCTR) and siRNA targeting SIRT1 or PGC-1α, then treated with HPF-CM or CAF-CM for 48 hours. **A-B.** PC3 cells silenced for SIRT1 were subjected to western blot (A) and invasion assay (B). **C-E**. PC3 cells silenced for PGC-1α and treated as in A were subjected to western blot (C), invasion assay (D) and TMRE analysis (E). Scale bar, 5 µm. TMRE fluorescence was quantified as signal measured per cell (E). Data represent mean ± SEM, *n =* 3. ns, no significant, *P < 0.05; **P < 0.005; ***P < 0.001, by two-tailed *t* test. **F-G.** Measurement of OCR and TCA cycle intermediates in PCa cells silenced for SIRT1 and exposed to HPF-CM or CAF-CM for 48 hours. Quantification analysis of protein levels in WB was relative to their respective control (the fold change numbers were indicated).

### CAF-induced TCA cycle deregulation affects HIF-1α activation, sustaining PCa cells invasion

Succinate and fumarate accumulation has been already reported to competitively inhibit α-KG-dependent prolyl hydroxylases (PHD), thereby stabilizing HIF-1α in SDH- and FH-deficient cells [20]. Similarly, among the electron transport chain (ETC) components, we found decreased succinate dehydrogenase (SDH) expression (i.e. complex II, **Fig. 4A**) and enzymatic activity (**Fig. 4B**) in CAF-treated PC3 cells. Interestingly, the downregulation in Complexes II-III, concomitant with Complex I increased expression (**Fig. 4A**), which are involved in redox homeostasis and mitochondrial reactive oxygen species (mtROS) generation, suggests that CAFs could promote ETC dysfunction and subsequent mitochondrial metabolic reprogramming in PC3 cells.

**Figure 4.**
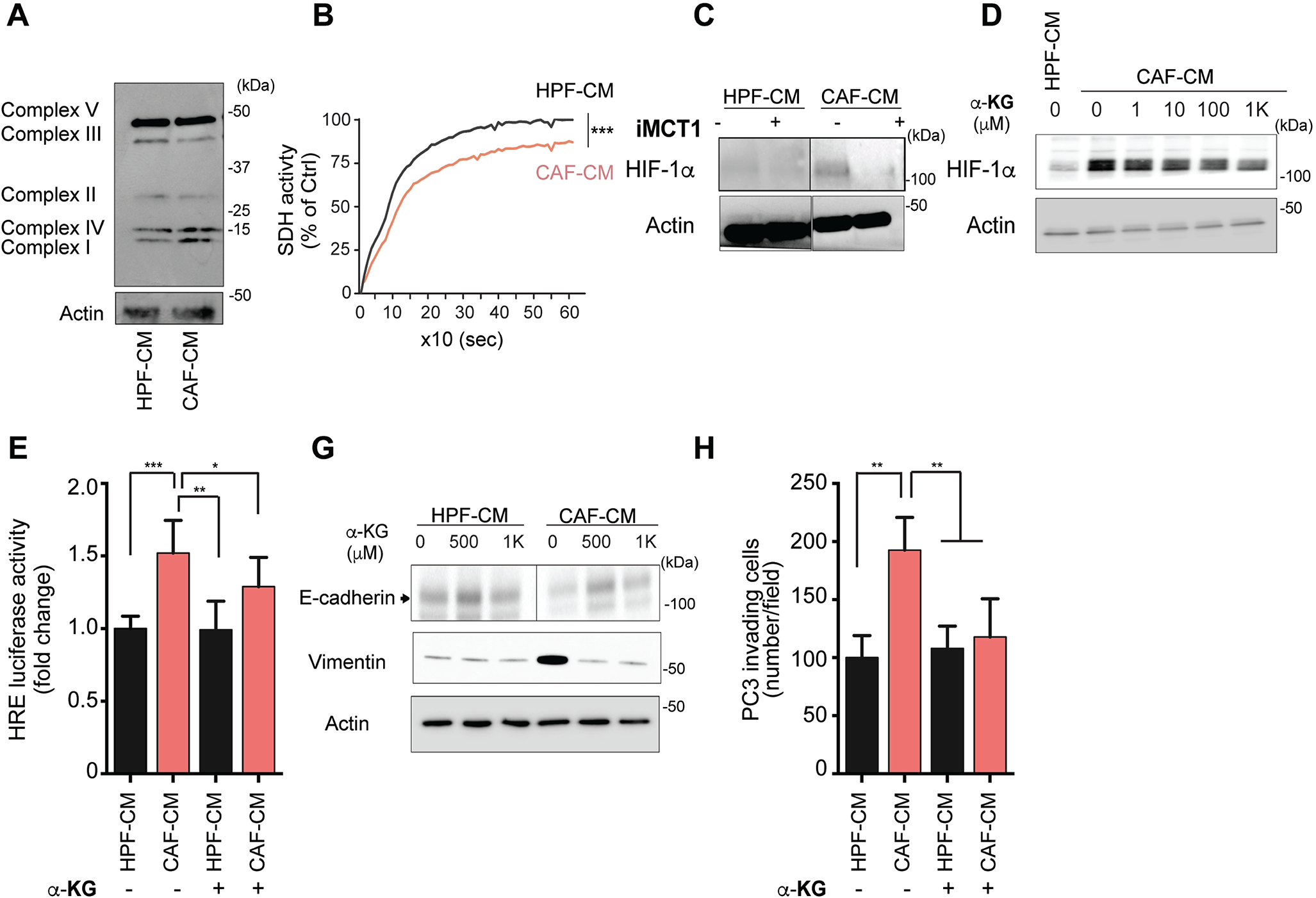
CAFs-promoted TCA cycle intermediates accumulation affects HIF-1-mediated invasive features. **A-B**. Analysis of ETC complexes expression by WB (A) and SDH activity (B) by enzymatic assay, in PC3 cells exposed to HPF-or CAF-CM for 48 hours. Mean ± SEM, *n =* 3. ***P < 0.001, by two-tailed *t* test. **D.** Analysis of HIF-1α expression in 48 hours CAF-CM-exposed PC3 cells upon impairment of lactate import through MCT1 inhibitor treatment. **E-F**. HIF-1α expression and activity were respectively determined by WB and HRE luciferase assay in PC3 cells treated with α-KG at the indicated concentration for 48 hours. **G-H.** E-cadherin levels and cell invasion were measured in α-KG-treated (1mM) PC3 cells. Data represent mean ± SEM, ns, no significant, p>0.05, *P < 0.05; **P < 0.005; ***P < 0.001, by one-way ANOVA, with Tukey post hoc tests.

We previously reported that CAF-induced HIF-1α stabilization and activation in PC3 cells occurs in normoxic condition [19]. Thus, we investigated whether CAF-driven lactate entry and its mitochondrial exploitation (*via* succinate accumulation) could be involved in promoting HIF-1 levels through pyruvate/succinate competitive actions linked to PHDs inhibition [21]. Crucially, inhibiting lactate import by using the MCT1 inhibitor reverted CAF-CM-induced HIF-1α stabilization in cancer cells (**Fig. 4C**). Similarly, administration of α-KG as succinate competitor prevented CAF-CM-induced HIF-1α stabilization and transcriptional activation in PC3 cells (**Fig. 4D-E**), ultimately impairing CAF-induced EMT and invasion (**Fig. 4G-H**).

### mtROS generation sustains the pro-invasive features induced by CAF exposure in PCa cells

Dysfunctional mitochondrial activity and/or high exploitation of mitochondrial respiration lead to enhanced mtROS accumulation that regulate cellular proliferation, migration and invasion [22–24]. As detected by FACS and confocal analysis, both PC3 and DU145 cells exposed to CAF-CM showed a significant increase in mtROS (**Fig. 5A and Fig. S2A**). Notably, mitoTEMPO administration, a specific scavenger of mitochondrial superoxide [25, 26], was able to interfere with CAF-induced EMT (**Fig. S2B**) and invasive abilities of PCa cells (**Fig. 5B**), thus suggesting mtROS as key drivers of the observed CAF-induced tumor cell aggressiveness.

**Figure 5.**
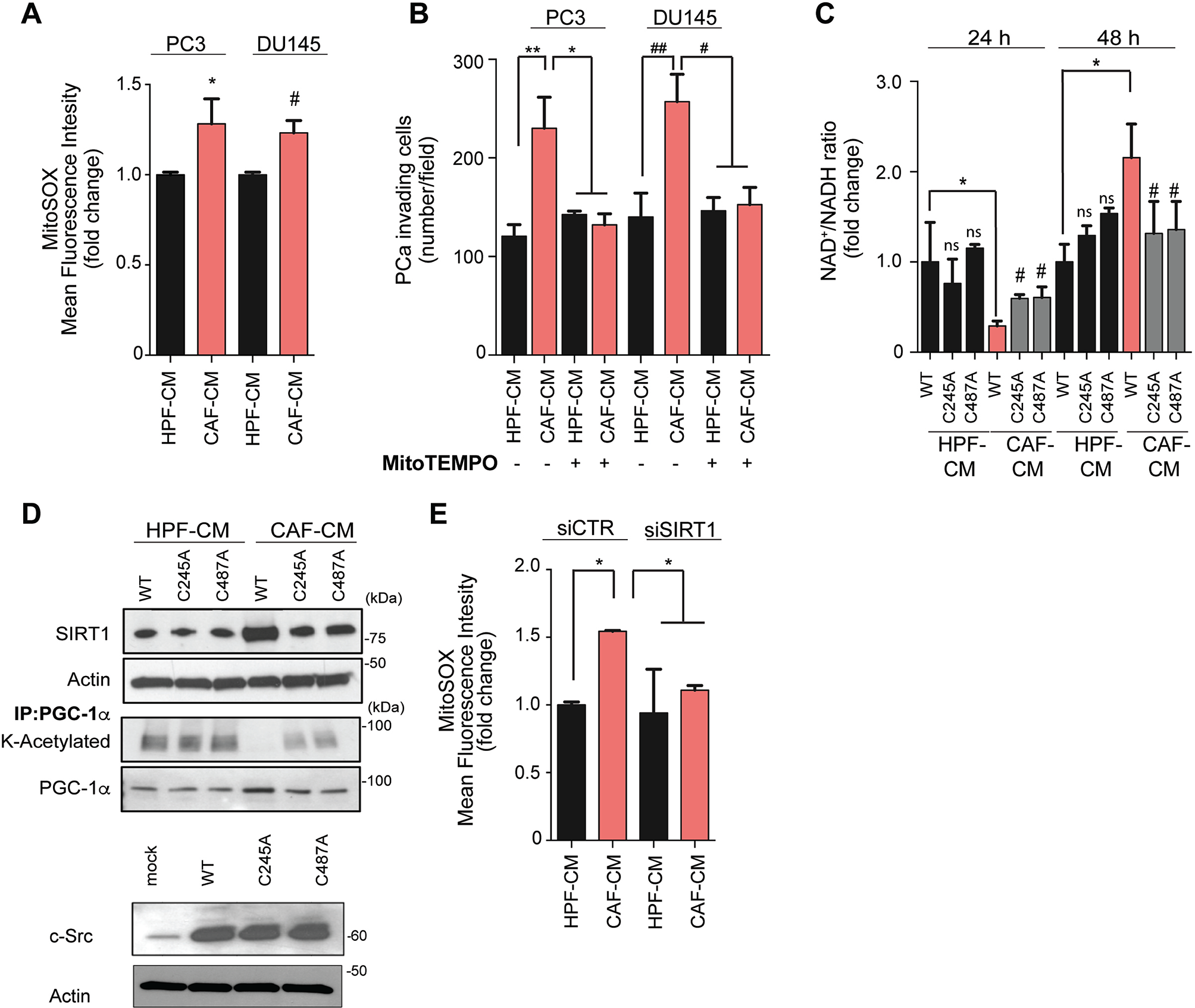
CAFs contact promotes mtROS accumulation sustaining motile features and SIRT1/PGC-1α activation in PCa cells. **A**. Evaluation of mtROS content by FACS analysis in PC3 and DU145 cells, after 48 hours treatment with HPF-or CAF-CM. **B.** PC3 cell invasion performed with or without MitoTEMPO administration (50 µM). **C-D** Evaluation of NAD^+^/NADH ratio (C) and PGC1-α acetylation status (D) in redox-insensitive Src mutants-transfected PC3 cells treated as in A. Mean ± SEM, *n* = 3. ns, no significant, p>0.05; *P < 0.05, by one-way ANOVA, with Tukey post hoc tests. **E.** Evaluation of mtROS content in PC3 cells silenced for SIRT1 and exposed to HPF-or CAF-CM. Data represent mean ± SEM, *n =* 3. ns, no significant, p>0.05; *P < 0.05; **P < 0.005, by two-tailed *t* test.

mtROS induce pro-migratory signals and metastasis via different pathways activation [24, 27] including that of the redox-sensitive tyrosine kinase Src [24]. Moreover, we have previously reported that Src-dependent PKM2 phosphorylation drives PCa cells reprogramming upon CAF-conditioning [3, 28]. Therefore, we analyzed Src and PKM2 redox state after CAF-conditioning, in the presence or absence of mitoTEMPO. Our results indicate that both enzymes undergo a strong oxidation after CAF exposure, which was prevented by mitoTEMPO (**Fig. S2C**). As Src oxidation (*i.e.* activation) has been reported in supporting CAF-induced lactate-addiction of PCa cells [3], we approached SIRT1/PGC-1α pathway by using two redox-insensitive Src mutants (Src C245A and C487A) [28]. Both non-oxidable Src mutants effectively impaired the lactate-dependent NADH/NAD^+^ unbalance observed upon CAF-CM administration in PC3 cells (**Fig. 5C and S2D**). The non-oxidable Src mutants also impairs the CAF-induced SIRT1 increased expression and activity, monitored by PGC-1α deacetylation (**Fig. 5D**). Similarly, SIRT1 silencing impaired CAF-induced mtROS generation in PC3 cells, possibly altering cancer cells ability to sense the CAF-released lactate (**Fig. 5E**).

### CAFs transfer mitochondria to PCa cells further enhancing their metabolic and motile features

We next analyzed the possibility that stroma influence mitochondrial/metabolic tumor cell behavior through an exogenous way, that is an horizontal transfer of mitochondria. For this goal, CAFs mitochondria were transiently labeled with MitoTracker Green and incubated with unlabeled PC3 or DU145 cells. Confocal analysis of coculture showed that PCa cells became positive for mitochondrial staining, suggesting the acquisition of stromal-derived mitochondria (**Fig. 6A-B**). The ability of mitochondrial transfer is a specific feature of the cancer-reprogrammed highly glycolytic CAFs [2], since the healthy counterpart (HPFs) are not able to donate their own mitochondria to PCa cells (**Fig. S3A**). In addition, no transfer of stained mitochondria was revealed from PC3 towards unlabeled CAFs, highlighting a specific unidirectional transfer of mitochondria from CAFs to PC3 cells (data not shown).

**Figure 6.**
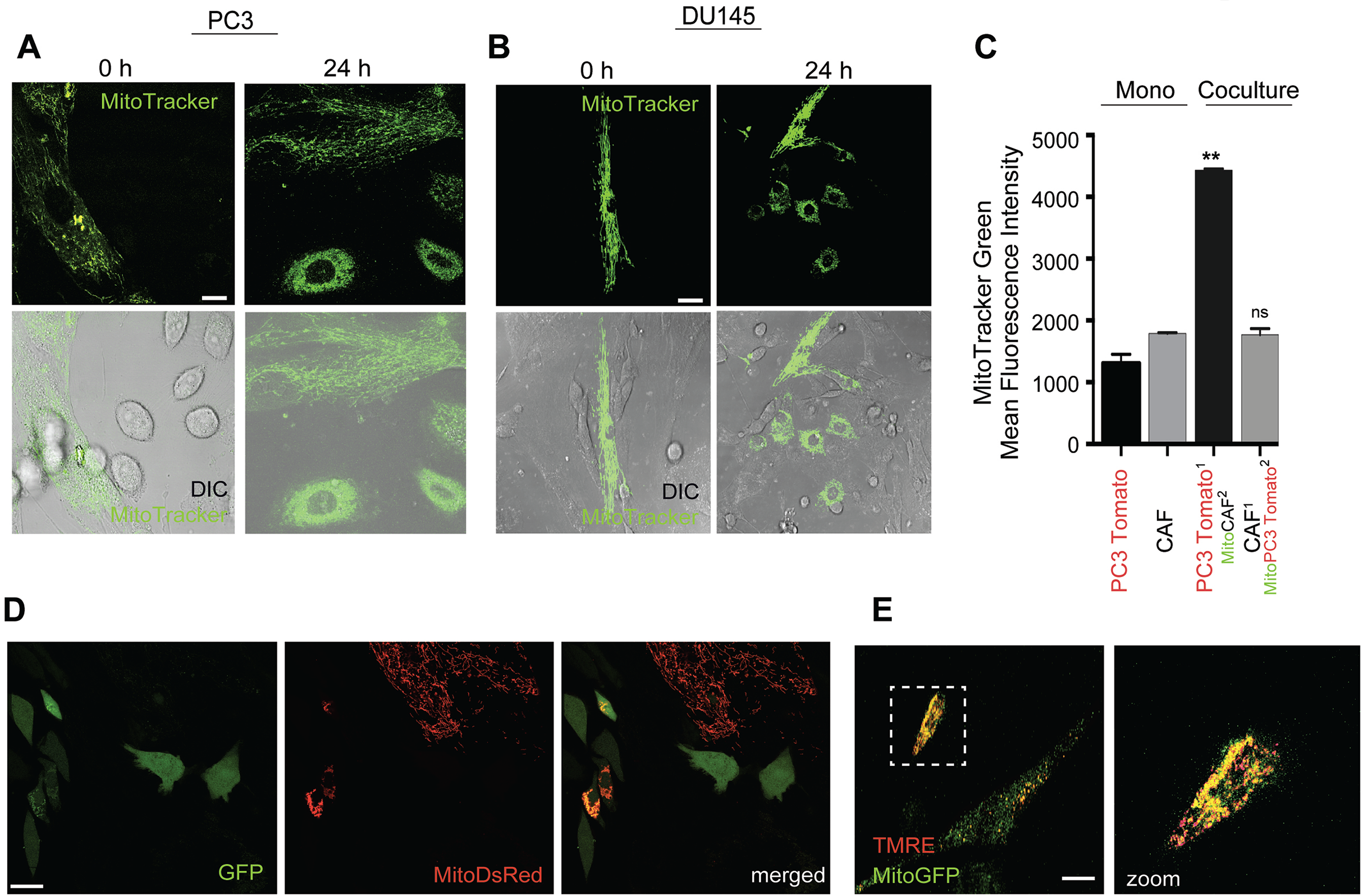
CAFs mediate horizontal transfer of mitochondria to PCa cells. **A-B**. CAFs-to-PCa cells mitochondria transfer analyzed by confocal microscopy. The transfer was quantified through the acquisition of MitoTracker Green labelling in unlabeled PC3 (A) or DU145 cells (B) after a 24 hours coculture with MitoTracker Green-stained CAFs. **C**. FACS analysis of acceptor cell populations (^1^) acquiring MitoTracker Green staining from donor cell population (^2^) in co-culture between CAFs and TdTomato fluorescent PC3 cells. Monoculture was used as comparator. Mean ± SEM in (C), *n* = 3. ns, no significant, **P < 0.05, by one-way ANOVA, with Tukey post hoc tests. Scale bar, 10 µm. **D.** Representative images of mitochondria transfer from CAFs transfected with mitochondria-targeted plasmids (MitoDsRed) to GFP-tracked PC3 cells after 24 h of co-colture. Scale bar, 10 µm. **E.** Representative images of TMRE staining upon 24 hours of MitoGFP-tracked CAF:unlabeled PC3 cells of co-culture. Scale bar, 10 µm; dashed lines indicate selected area for 63X zoom of TMRE/MitoGFP-loaded PC3 cell.

CAF mitochondria transfer to PC3 cells was also confirmed in coculture between differentially labeled cells (PC3 cells stably expressing the red fluorescent protein tdTomato-PC3Tom, and CAFs with MitoTracker Green-prelabelled mitochondria). FACS analysis revealed a red and green double staining of PC3 cells, confirming the transfer of green-fluorescent mitochondria to red-fluorescent PC3 cells (**Fig. 6C**). To exclude that the staining of PC3 cells was due to dye leakage from CAFs, we alternatively established cocultures between GFP-transfected PC3 cells and CAFs transfected with a plasmid encoding a mitochondrial-targeted fluorescent protein (MitoDsRed). Indeed, we confirmed that MitoDsRed stained-mitochondria are proficiently transferred from CAFs to GFP-labelled PC3 cells (**Fig. 6D**). Notably, we also demonstrated that both endogenous tumor cell mitochondria and exogenously acquired mitochondria from MitoGFP-labelled CAFs have a functionality-associated fluorescent signal (**Fig. 6E**), suggesting that CAFs increase PCa cell respiratory capacity not only by enhancing the activity of cancer cell endogenous mitochondria (*via* the SIRT1/PGC-1α pathway), but also by recruiting functional mitochondria from symbiotic stromal cells.

To investigate whether PCa cells *in vivo* recruited mitochondria from CAFs, we established tumor xenografts in immunodeficient SCID-bg/bg mice by subcutaneously injecting either unlabeled or PC3Tom cells together with unlabeled or MitoGFP-expressing CAFs, respectively (**Fig. 7A**). After 25 days, tumors were excised and disaggregated for cell recovery. Tumors from different groups were weighed after excision to ensure that the expression of fluorescent proteins did not interfere with cell growth *in vivo*. To detect the *in vivo* transfer of MitoGFP-labelled mitochondria from CAF donor cells to PC3 receiving cells, PC3Tom cells recovered by tumors excised by animals single-injected with PC3Tom or co-injected with PC3Tom + mitoGFP-CAFs, were analyzed by FACS and confocal microscopy. A significant enrichment in mitoGFP-stained PC3Tom cells, due to the acquisition of CAF-released mitoGFP-labelled mitochondria (**Fig. 7B-C**) was revealed, highlighting the importance of the unidirectional mitochondrial transfer *in vivo*. Furthermore, by analyzing PC3 cells isolated from tumors excised by animals injected with unlabeled cells (PC3 single injection or PC3 + CAF co-injection), we observed an increase in both the overall mitochondrial content, measured by MitoTracker Green staining, (**Fig. 7D-E**) and functionality, analyzed by TMRE staining (**Fig. 7F**), upon *in vivo* CAF interaction. These metabolic data were also supported by the higher expression of MCT1 and SIRT1 and the downregulation of E-cadherin, observed in PC3 cells *in vivo* interacting with CAFs with respect to PC3 cells alone (**Fig. 7G**), in keeping with their lactate exploitation, OXPHOS addiction and EMT execution.

**Figure 7.**
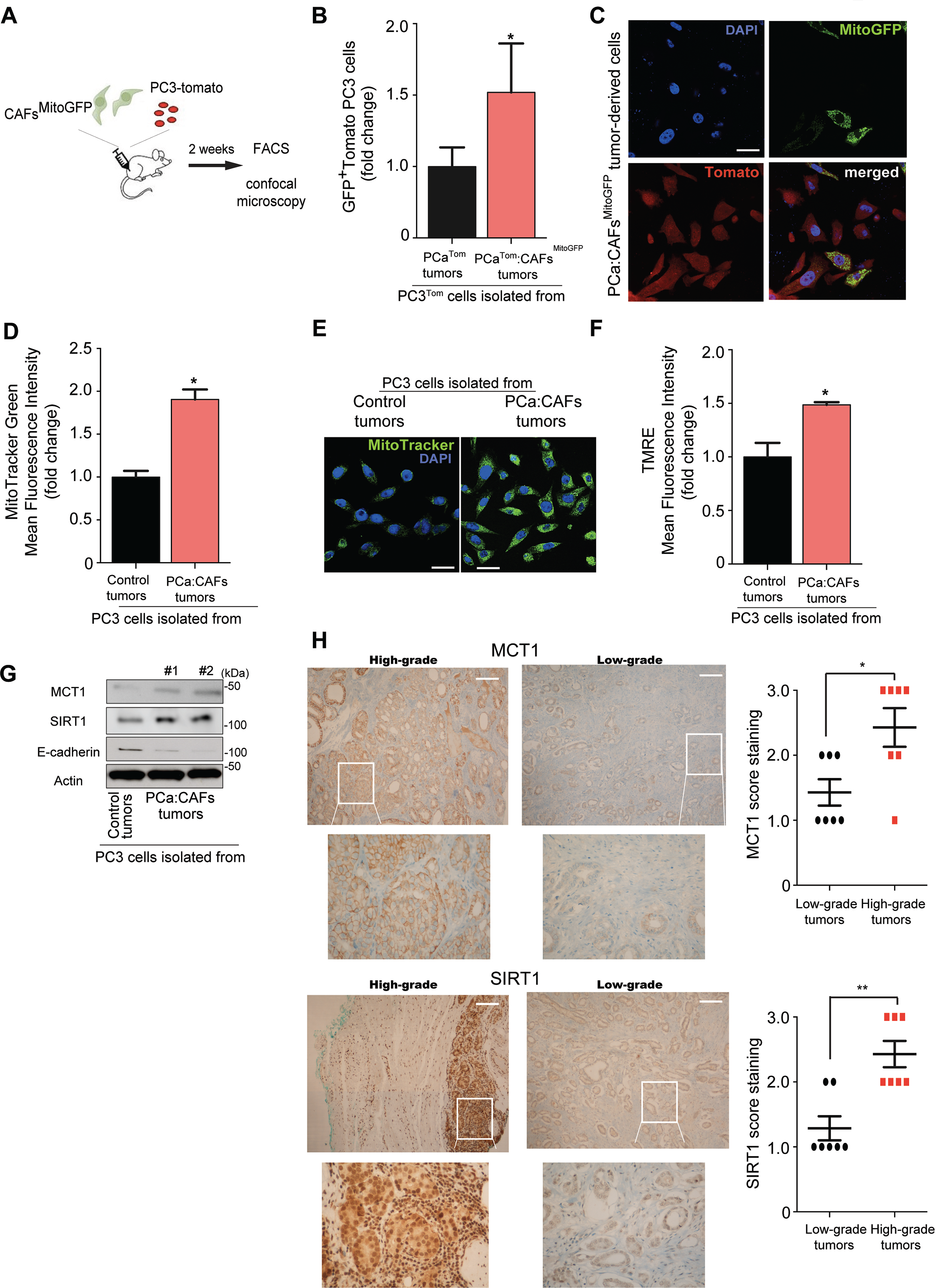
*In vivo* transfer of mitochondria from CAFs to PCa cells. **A.** Schematic model of the experimental setting. **B-C.** FACS and confocal analysis showing the acquisition of labelled mitochondria derived from MitoGFP-infected CAFs by TdTomato-fluorescent PC3 cells recovered after tumor disaggregation. DAPI was used to stain nuclei. Scale bar, 10 µm. **D-F.** Mitochondrial mass and mitochondrial membrane potential were measured in PC3 cells derived from disaggregated tumors by FACS or confocal analysis upon MitoTracker Green or TMRE staining, respectively. Scale bar, 10 µm. **G.** Expression of MCT1, SIRT1 and E-cadherin in PC3 cells derived from disaggregated tumors (#1 and #2 are relative to tumors excised from two different mice). Data represent mean ± SEM, *n =* 3. **P < 0.05; **P < 0.005, by two-tailed *t* test. **H.** Representative IHC analysis of MCT1 and SIRT1 in low-grade (*grade group 1*) and high-grade (*grade group 3*) prostate tumors and relative quantification per intensity of staining scoring. Data shown are from 7 samples per group ± SEM, Student *t* test. Scale bar, 200 μm. Magnification (40X) of highlighted areas was shown below.

In order to evaluate whether *in vivo* higher grade of malignancy correlates with an increase in lactate exploitation, we analyzed explants from patients affected by clinically localized well differentiated adenocarcinoma (grade group 1) or low differentiated PCa with extension in the seminal vesicle and metastasis in the lymph-nodes and low differentiated adenocarcinomas (grade group 3). We observed a significantly higher expression of MCT1 and SIRT1 high grade tumors, while low levels of both the proteins are expressed in the low grade tumors (Fig. 7H). These *in vivo* evidence reinforce our conclusion about the role of MCT1-dependent lactate influx and the activation of the SIRT1-mediated enhancement of mitochondrial metabolism in conferring more malignant traits, likely contributing to the improvement of metastatic potential.

It has been reported that mitochondrial transfer can occur by endocytosis, exosomes [29] or *via* cytoplasmic bridges, also known as tunneling nanotubes [30], that allow the transfer of mitochondria from adjacent cells [31–33]. Intriguingly, confocal images obtained during co-culture of PC3 cells with MitoGFP-transfected CAFs indicated the formation of physical connection between stromal and tumor cells with labelled mitochondria located within these cellular bridges, leading to the transfer of mitochondria from CAFs to PCa cells (**Fig. 8A**-**B**). To evaluate the impact of stromal transferred mitochondria on PC3 cells, green fluorescent mitochondria from CAFs were isolated, and administrated to tumor cells, upon confirming their functionality by OCR analysis (data not shown) (**Fig. 8C**). Representative images showed that PC3 cells significantly upload exogenous mitochondria, whereas CAFs are not able to act as recipient cells, suggesting a characteristic feature of PC3 cells to acquire mitochondria from outside (**Fig. 8D, left**). We also observed that PC3 cells exposed to CAF-CM are more prone to incorporate isolated mitochondria, as also reported by the quantification analysis (**Fig. 8D, right**). Particularly, z-stack-confocal imaging of a representative receiver cancer cell and the associated 3D reconstruction clearly demonstrated that the cancer cells (red fluorescent) have incorporated intact green-stained mitochondria (**Fig. 8E**, **Video 1**).

**Figure 8.**
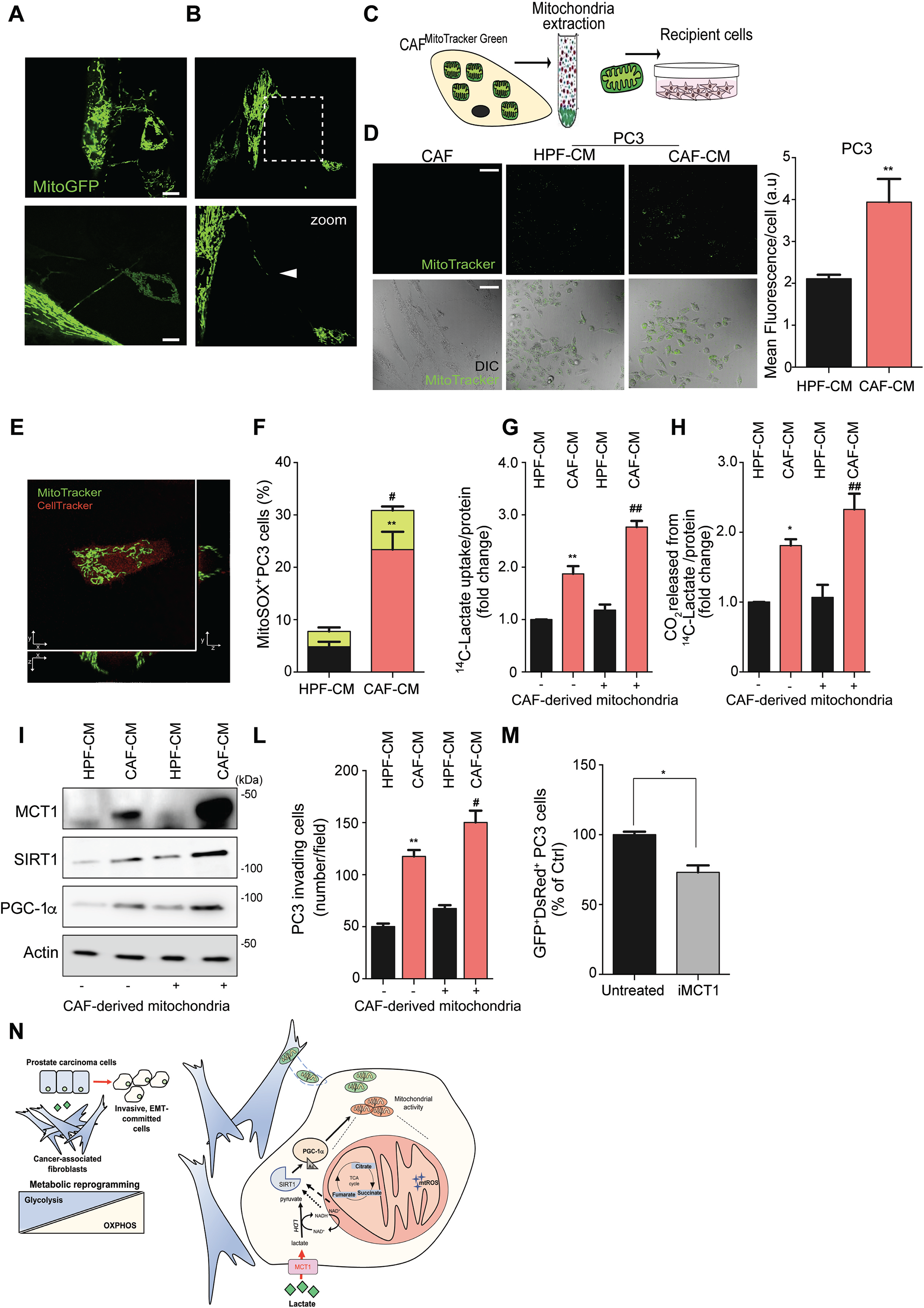
Transferred mitochondria from CAFs enhance motile features and lactate metabolism of PCa cells. **A-B.** Two representative images of tunnelling structures mediating mitochondria transfer from MitoGFP-transfected CAFs toward PC3 or DU145 cells. Scale bar, 10 µm. **C.** Schematic model for the 24 hours-administration of CAFs-isolated mitochondria to PC3 cells (previously treated with HPF-or CAF-CM for 48h) or to CAFs (1µg mitochondria/1.5×10^5^ cells). **D.** Fluorescence images showed MitoTracker Green dots as readout of mitochondria uptake in the indicated cell types (PC3, CAF) and treatments (HPF-or CAF-CM). Scale bar, 10 µm. Mean fluorescence inside cells was measured and plotted. **E.** Internalization of mitochondria was confirmed by confocal z sections in pre-labeled OrangeRed PC3 cells acquiring MitoTracker Green-labeled CAF mitochondria. Scale bar, 10 µm. **F.** Evaluation of mtROS by FACS analysis in PC3 cells treated as above; green bars are representative for the percentage of PC3 cells acquiring CAF-derived MitoTracker Green-labeled mitochondria. Mean ± SEM, *n* = 3. **P < 0.05; **P < 0.005, by two-tailed *t* test. **G-H.** Evaluation of lactate uptake (G) and CO_2_ release (H) by using radiolabeled ^14^C-(U)-lactate in PC3 cells, exposed to HPF-or CAF-CM (48 hours), upon CAF mitochondria administration (24 hours). Mean ± SEM, *n* = 3. **P < 0.05; **P < 0.005, by two-tailed *t* test. **I-L.** Western blot analysis for MCT1, SIRT1 and PGC-1α expression (I) and invasion assay (L), performed on PC3 cells, treated for 48 hours with HPF-or CAF-CM and subsequently exposed to CAF mitochondria for 24 hours. Mean ± SEM, *n* = 3. **P < 0.05; **P < 0.005, by two-tailed *t* test. **M.** FACS analysis showing the inhibition of mitochondria transfer from MitoDsRed-transfected CAFs to GFP expressing-PC3 cells, upon impairment of lactate influx by 24 hours MCT1 inhibitor (AR-C155858) treatment (10µM). Data represent mean ± SEM, *n =* 3. **P < 0.05; **P < 0.005; #P< 0.05; ##P<0.005 between mitochondria-loading PC3 cell populations, by two-tailed *t* test. **N**. Graphical scheme depicting the model of the prostate tumor-stroma metabolic interaction. CAFs potentiates PCa cell mitochondria by activating the lactate/SIRT1/PGC1-α axis and by supplying their own functional mitochondria to boost the oxidative metabolism of PCa cells. The subsequent OXPHOS engagement of PCa cells is associated to EMT and to a more invasive phenotype.

We next investigated whether the observed mitochondrial transfer impacts on CAF-dependent PCa cell phenotype and aggressive features. Indeed, we exposed cancer cells to mitochondria isolated from MitoGFP-transfected CAFs and we measured mtROS production upon CAF-CM or HPF-CM exposure (**Fig. 8F**). As previously shown, mtROS production is enhanced in CAF-CM treated PC3 cells (**Fig. 8F, black vs red bars**), but, interestingly, the fraction of cells acquiring CAF mitochondria (MitoGFP^+^) displayed a further enhancement in MitoSOX staining, compared to MitoGFP^-^ cells (**Fig. 8F, green bars**). We hypothesized that tumor cell avidity for exogenous CAF-derived mitochondria could be associated to a higher lactate exploitation. Notably, PC3 cells that have been reprogrammed by CAF-CM administration, increased their lactate uptake (**Fig. 8G**) and the lactate OXPHOS-dependent utilization, assayed by radioactive ^14^CO_2_ tracing (**Fig. 8H**), after the addition of CAF-derived mitochondria. This metabolic reprogramming exerted by CAF-derived mitochondria supplementation leads to a higher activation of the described lactate-dependent SIRT1-PGC1α axis: indeed, PC3 cells showed higher expression of MCT1, SIRT1 and PGC-1α levels (**Fig. 8I**) when exposed to CAF-derived mitochondria, thereby resulting in a significant increase in stroma-potentiated tumor invasive abilities (**Fig. 8L**). Crucially, we further evidenced that such stroma-to-tumor mitochondrial transfer is impaired by hindering lactate-dependent tumor:CAFs crosstalk by MCT1 inhibition (**Fig. 8M**).

## DISCUSSION

Metabolic reprogramming is essential to satisfy the different cell requirements during tumor initiation and progression. Notably, such reprogramming is not merely influenced by cell-autonomous genomic alterations but also by non-genomic components and non-cell autonomous players, e.g. the tumor microenvironment. According to the so-called “two compartment” model of tumor metabolism [34], anabolic malignant cells can establish a metabolic symbiosis with the adjacent stromal catabolic cells to subtract high-energy metabolites (e.g. lactate, glutamine, fatty acids) that can stimulate tumor proliferation and foster metastatic spreading [35]. Particularly, CAFs-derived lactate is the main driver for the enhancement of PCa cells malignancy, achieved by sustaining a complex metabolic and molecular rearrangement involving PKM2/miR-205/EMT partnership [3].

Since OXPHOS metabolism was crucial to exploit the energy rich metabolites released by highly glycolytic stromal cells, we focused our attention on the mitochondrial rewiring that prostate carcinoma cells could undergo when in contact with the CAF secretome.

Lactate released from CAF generates an unbalance in the NADH/NAD^+^ ratio of the PCa cells (**Fig. 1**). While NADH early increase could be due to the reversible activity (lactate→pyruvate) and mitochondrial localization of the lactate dehydrogenase (LDH) [36], NAD^+^ subsequent high levels respond to the subsequent mitochondrial activation of stromal reprogrammed PCa cells, resulting in the SIRT1-mediated PGC-1α activation (a schematic model has been proposed in **Fig. 8N**). SIRT1 is a crucial “cog” in the molecular machine driving CAF-induced mitochondrial re-education of PCa cells. Indeed, the impairment of SIRT1 function by a genetic approach counteracts EMT execution and invasiveness, through a clear impact on HIF-1α stabilization. PGC-1α has been reported to have a crucial role in circulating breast cancer cells [8] and is a major driver of metastatic localization of breast cancer cells to the brain [37] and to the lung [38]. However, other studies have reported an opposite role, showing an association of PGC-1α downregulation with disease progression in PCa [9]. Notably, our CAF-exposed PCa model relies on the SIRT1-dependent acetylation of PGC-1α and not exclusively on its expression, possibly explaining the different roles of PGC-1α reported in these studies. Indeed, we argue that PGC-1α most likely reprograms and creates a new metabolic state that can either promote or hinder tumor growth and metastatic progression, depending on metabolic environment that exists within the primary and metastatic sites (i.e. circulating lactate). It is conceivable that when a cancer cell leaves the primary site to colonize secondary sites, PGC-1α deacetylation (i.e. activation) coordinates the bioenergetic profile of the circulating cancer cell, particularly reprogramming the mitochondrial status and activity, in order to have a high adaptability and plasticity for high-energy nutrients exploitation and ATP production to survive (when circulating) allowing to engraft in different secondary sites metabolic environment [39].

In the model used in the current study, mtROS production and handling links the molecular rearrangements induced by CAF-conditioning to the metabolic reprogramming and finally to the phenotypical changes observed. Indeed, mtROS are established modulators of the metastatic cascade [24] and our data clearly showed a CAFs-mediated mtROS generation, responsible for the oxidation of molecular mediators, i.e. Src and PKM2, crucial for sustaining the metabolic circuitry and EMT engagement in CAFs-exposed PCa cells. The concomitant mtROS generation and PGC-1α activation, a known driver of anti-oxidant response, seems controversial. However, PGC-1α could control redox homeostasis *via* different pathways. Indeed, mtROS triggers AMPK-PGC-1α signaling as feedback loop to limit potentially damaging levels of ROS [40]. Although we did not investigate NADPH levels and anti-oxidant response in our model, we can speculate that finely regulated mtROS levels could act as signaling molecules in the metabolic crosstalk between CAFs and PCa cells. Interestingly, mtROS scavenging is able to prevent PCa cells malignant rewiring and this may partially explain why mitochondria-targeted antioxidant drugs are effective *in vivo* to inhibit tumor metastasis [7, 24, 26], whereas broad-range antioxidants (e.g. N-acetyl-cysteine) tend to fail probably due to their inability to target and lower ROS generated and localized within the mitochondria [41].

Lactate-dependent enhancement of mitochondrial function also influences TCA cycle intermediates, mainly increasing citrate, malate, succinate and fumarate. Such activation of OXPHOS metabolism in cancer cells in a lactate-dependent manner was recently supported by a study showing that circulating lactate can be used by tumor cells to fuel the TCA cycle [42]. The CAF-mediated increase succinate levels can be explained either by a TCA cycle overload and/or by a SDH activity dysfunction. We confirmed that CAF-CM impaired SDH expression and activity in PCa cells and leads to succinate accumulation, resulting in HIF-1α stabilization/activation and EMT execution, as reported in other models [43]. Similarly to succinate, fumarate promotes EMT signature in cancer cells, via epigenetic downregulation of miR-200ba429 [44].

Besides the reported enhancement and reshaping of endogenous mitochondria upon CAF-CM exposure, for the first time we provide evidence about the ability of PCa cells to receive mitochondria from neighboring stromal cells, even in the absence of mitochondrial defects (i.e., mtDNA mutations).

Growing evidence showed that reprogrammed stromal cells, undergoing glycolysis, display mitochondrial dysfunction and activate autophagy/mitophagy, which results in a series of stimuli ultimately leading to enhancement of the mitochondrial function of adjacent epithelial cancer cells [45, 46]. Here, we hypothesized that highly glycolytic CAFs could donate their dispensable mitochondria to neighboring PCa cells to improve their malignancy, concurring to strengthen the effect of soluble factors [19, 47] and/or exosomes [48]. According to our data, only few PCa cells are able to act as “receivers” for exogenous mitochondria. We hypothesize that during co-colture, paracrine signals govern the establishment of a “chemotactic-like force” generated by the receiver cells in guiding the tunneling nanotube formation and the unidirectional transfer of mitochondria. When CAF-derived mitochondria are administrated in the absence of CAFs:cancer cells contact, it is plausible that only few cells have been stimulated to act as “receiver” cells, hence only few cells could uptake exogenous mitochondria. In agreement with the idea that soluble secreted factors (e.g. CAF-released lactate) could influence cancer cells avidity towards external mitochondria, CAF-CM pretreatment of cancer cells prior to CAF-derived mitochondria administration enhances the receiving capacity of cancer cells. We speculate that without this priming event, cancer cells are not prone to acquire exogenous mitochondria, but when they are primed to act as “receivers”, they can potentially accept mitochondria independent on the “donors”. It is likely that within the prostate tumor microenvironment, only the cancer-reprogrammed highly glycolytic CAFs can act as “donor” cells of their dispensable mitochondria. On the contrary HPF, which still rely on a higher extent on mitochondrial metabolism, are not prone to deprive themselves of their mitochondria.

Notably, CAFs-mediated mitochondrial transfer phenotypically acts on CAFs-reprogrammed PCa cells, presumably by selecting a subpopulation of cells characterized by enhanced invasive abilities, especially inclined to import and exploit CAF-secreted lactate, as reported by the MCT1 increased expression and lactate utilization in mitochondria-enriched PCa cells. It may be conceivable that the SIRT1/ PGC-1α pathway can be involved in mediating the increase in lactate upload in mitochondria receiver cells. It has been reported that in skeletal muscle cells, PGC-1α upregulates lactate uptake by increasing the expression of MCT1 [49]. In addition recent evidence reported that PGC-1α enhances the expression and activation of lactate dehydrogenase B (LDH-B), which drives the conversion of lactate to pyruvate, while repressing the LDH-A activity, thus resulting in the enhancement of lactate consumption [50]. Independently on the *primum movens*, whether this transfer of mitochondria is a rare event or a highly impacting phenomenon to improve cancer malignancy and whether this process has clinical relevance is still unknown and require additional investigation.

Taken together, our data suggest that the CAF-derived secretome confers to PCa cells a previously unexplored function, driving the activation of a SIRT1/PGC-1α axis that is responsible for enhancement of mitochondrial function and, in turn, for mtROS generation and oncometabolite accumulation. All these events concur to enhance the migratory and metastatic abilities of PCa cells. Particularly, our findings reveal that glycolytic CAFs directly transfer their functional but underexploited mitochondria to cancer cells, further promoting mitochondrial utilization and OXPHOS-addiction of PCa cells, and ultimately promoting their malignancy.

## MATERIALS AND METHODS

### Cell lines, antibodies and reagents

Human PCa cells (PC3, DU145) were obtained and authenticated by PCR/short tandem repeat (STR) analysis from ATCC and maintained at 37°C/5% CO_2_ in DMEM supplemented with 10% fetal bovine serum (FBS), 2 mM L-Glutamine and 1% penicillin/streptomycin. Human prostate fibroblasts (HPFs and CAFs) were isolated from surgical explants after patient informed consent, according to the Ethics Committee of the Azienda Ospedaliera Universitaria Careggi (Florence, Italy). Unless specified otherwise, all reagents were obtained from Sigma-Aldrich. The following antibodies were used in this study: rabbit E-Cadherin (#5296) and rabbit PKM2 (#4053) from Cell Signaling Technology, mouse HIF-1α (#610958) from BD Biosciences, rabbit total OXPHOS cocktail (ab110411) and rabbit acetyl lysine (ab80178) from Abcam, mouse SIRT1 (sc-74504), mouse MCT1 (sc-365501), rabbit PGC-1α (sc-13067), mouse vimentin (sc-6260), rabbit c-Src (sc-19) and mouse b-actin (sc-58673) from Santa Cruz Biotechnology, Inc. Secondary antibodies were from Santa Cruz Biotechnology, Inc. MitoTEMPO (sc-221945) was from Santa Cruz Biotechnology, Inc and MCT1 inhibitor AR-C155858 (#4960) was from Tocris BioScience.

### Conditioned media from fibroblasts

HPFs and CAFs were grown to subconfluence and treated for 48 hours with serum-free medium to obtain the corresponding CM. PCa cells were treated with CM collected from fibroblasts (HPFs or CAFs) for 24 or 48 hours.

### Plasmid and siRNA transfection

Cells were transfected with siSIRT1 and siPGC-1α or negative controls (Santa Cruz Biotechnology) or with mitochondria-targeted plasmids mt-HA-eGFP, AT-F001-D and mtDsRed, AT-F002-D (Aequotech srl) using Lipofectamine RNAi-Max Reagent or Lipofectamine 3000 (Thermo Fisher Scientific), respectively. Analyses were performed 3 days after transfection. For mitochondria tracking *in vivo*, CAFs were infected with lentiviral particles containing the pLJM1-EGFP plasmid.

### NAD^+^/NADH measurement

Intracellular NAD(H) content was quantified by means of an enzymatic cycling procedure according to [51].

### Measurement of oxygen consumption rate (OCR)

PCa cells (PC3, DU145) were maintained at 37 °C and oxygen consumption was measured using a Clark-type O_2_ electrode (Hansatech). The rate of decrease in oxygen content, related to cell number, was taken as index of the respiratory ability.

### Mitochondrial staining and mitochondrial reactive oxygen species (mtROS) analysis

MitoTracker Green, TMRE and MitoSOX were used accordingly to manufacturers’ instructions (Invitrogen). Cwll were analyzed by flow cytometry using FACScan flow cytometer (BD Biosciences) or by confocal imaging (Leica TCS SP5).

### SDH activity

Cell homogenates were incubated in a phosphate buffer containing sodium azide, 2,6 dichlorophenolindophenol (DCPIP), sodium succinate, and phenazine methosulfate. SDH specific activity was measured by photometry using the Victor3 1420 Multilabel Counter (Packard Instruments, Perkin-Elmer) to measure the decrease in absorbance that resulted from the oxidation of DCPIP at 600 nm.

### Radioactive assays

PCa cells were treated as described in the Figures and subjected to radioactive assays as described previously [2].

### Confocal image acquisition and analysis

Cells were stained with TMRE, DAPI or MitoTracker Green dye or transfected with fluorescence-tracked plasmids (mt-HA-eGFP,and mtDsRed, AT-F002-D). Fluorescence samples were examined using a microscope (TCS SP5; Leica) as described in detail in Supplementary Material and Methods. Fluorescence quantification and 3D reconstruction were assessed using ImageJ and Imaris Bitplane software, respectively.

### Mitochondria isolation and transfer to recipient cells

Mitochondria were prepared using a modified protocol described in [52]. For a detailed procedures, see Supplementary Material and Methods.

### Immunoprecipitation, immunoblotting and BIAM labeling

Cells were lysed in RIPA buffer and subjected to immunoprecipitation and/or Western blot as described previously [3]. Detection of protein redox state, was performed as described in [53]. Briefly, at the end-point of the experiment, PC3 cells were frozen rapidly in liquid nitrogen and exposed to oxygen-free RIPA Buffer containing 100 mM of N-(biotinoyl)-N0-(iodoacetyl) ethylenediamine (BIAM). After sonication, cell lysates were incubated for 15min at room temperature. Lysates were then clarified by centrifugation and 500 mg of each sample subjected to immunoprecipitation with c-Src or PKM2-specific antibodies. BIAM-labelled immunocomplexes were separated by SDS-PAGE and the biotinylated/reduced fraction of the immunoprecipitated proteins was revealed with Horseradish Peroxidase-conjugated streptavidin.

### In vitro Boyden invasion assay

Motility and invasion assays were conducted as described previously [3].

### TCA cycle metabolites quantification by HPLC-ESI-MS

PC3 cells (5 x 10^6^ cells/condition) were homogenized in water, added to methanol and chloroform, centrifuged and the aqueous layer was recovered. After addition of cold acetone, suspensions were centrifuged and supernatants used for HPLC-MS analysis using an LTQ Orbitrap mass spectrometer coupled to and Accela HPLC system (Thermo Fischer Scientific). For a detailed procedures, see Supplementary Material and Methods.

### Determination of PHD2 and HIF-1 activities

Dual luciferase reporter assays were performed with the dual luciferase kit (Promega) using pGL3-(PGK-HRE6)-TK-Luc as reporter of HIF-1 activity. Reporter plasmid and a Renilla transfection normalization vector (Promega) were used.

### Xenograft experiments

*In vivo* experiments were conducted in accordance with national guidelines and approved by the ethical committee of Animal Welfare Office of Italian Work Ministry and conform to the legal mandates and Italian guidelines for the care and maintenance of laboratory animals. Six-to 8-week-old male severe combined immunodeficient (SCID)-bg/bg mice (Charles River Laboratories International) were s.c. injected with 1 × 10^6^ PC3 cells or mixed with 0.5 × 10^6^ of fibroblasts for co-injection. For cell tracking and mitochondria transfer experiments, PC3 TdTomato and MitoGFP fibroblasts were used.

### Immunohistochemistry on tissue specimens

IHC was performed on 14 paraffin-embedded and serially cut prostate cancer tissues obtained from Azienda Ospedaliera Universitaria Careggi, Florence, following patient consent and approval from the local Ethics Board. For all the histological samples the Gleason Grade and the pathological T staging were defined. The cases included in the study were divided in two groups according to histologic features: 1) prostatic adenocarcinoma, acinar type, Gleason score 3+3=6, grade group 1, stage pT2, pN0; 2) prostatic adenocarcinoma, acinar type, Gleason score 4+3=7, pattern 4: 70-80%, grade group 3, stage pT3b, pN1. IHC staining was performed using the following antibodies: MCT1 (H-1), mouse monoclonal antibody (Santa Cruz Biotecnology) and SIRT1 (B-10), mouse monoclonal antiboby (Santa Cruz Biotecnology) on a Ventana BenchMark ULTRA immunostainer (Ventana Medical Systems, Tucson, AZ). For semiquantitative assessment of the IHC data, the mean percentage of positive tumor cells was determined at 400X magnification for each section and evaluated in a blinded manner. Sections were graded on the basis of the intensity of staining as 0 (negative), 1 (weak), 2 (moderate), or 3 (strong).

### Tumor dissociation and *ex vivo* culture

Briefly, mice were sacrificed when tumors reached 0.5 cm^3^. Excised tumors were minced into 2-4 mm fragments, which were then incubated for 2h with the dissociation solutions containing 200 U/ml collagenase III, IV and Hyaluronidase. Digested fragments were filtered (70 μm cell strainer) and released cells were freshly analyzed.

### Statistical analysis

Statistical analysis was performed with Prism software (GraphPad Software). Data show means +/-SEM from at least three independent biological replicates (indicated as “*n*”). Comparisons between groups were analyzed by t-test or one-way/two-way analysis of variance (ANOVA) followed by post-hoc test. Statistical significance was considered when p<0.05.

## ACKNOWLEDGMENTS

The work was supported by Fondazione Umberto Veronesi to AM, Associazione Italiana Ricerca sul Cancro (AIRC) (grant 8797 to PC), AIRC and Fondazione Cassa di Risparmio di Firenze (grant 19515 to PC and AM), Istituto Toscano Tumori (grant 0203607 to PC), Programma operativo regionale Obiettivo “Competitività regionale e occupazione” della Regione Toscana cofinanziato dal Fondo europeo di sviluppo regionale 2007-2013 (POR CReO FESR 2007-2013, grant to PC), Interuniversity Attraction Pole from Belspo (grant #UP7-03 to PS), an Action de Recherche Concertée from the Communauté Française de Belgique (ARC 14/19-058 to PS), and the Belgian Fonds National de la Recherche Scientifique (F.R.S.-FNRS, to PS).

The authors thank: Dr Paolo E. Porporato (University of Turin, Italy) for providing mitochondria-targeted plasmids mt-HA-eGFP, AT-F001-D and mtDsRed, AT-F002-D (Aequotech srl); Dr Andrea Rasola (University of Padua, Italy) for providing pLJM1-EGFP plasmid for lentiviral infection; Dr Barbara Stecca (Istituto Toscano Tumori, Florence, Italy) for providing pCMV-dR8.91 packaging plasmid and pMD2G envelope plasmid; Dr. Nicla Lorito and Dr. Lavinia Ferrone for technical support. PS is a F.R.S.-FNRS Senior Research Associate. The MASSMET platform (https://www.uclouvain.be/en-massmet.html) is acknowledged for the access to the HLPC-MS.

---

**Figure S1.**
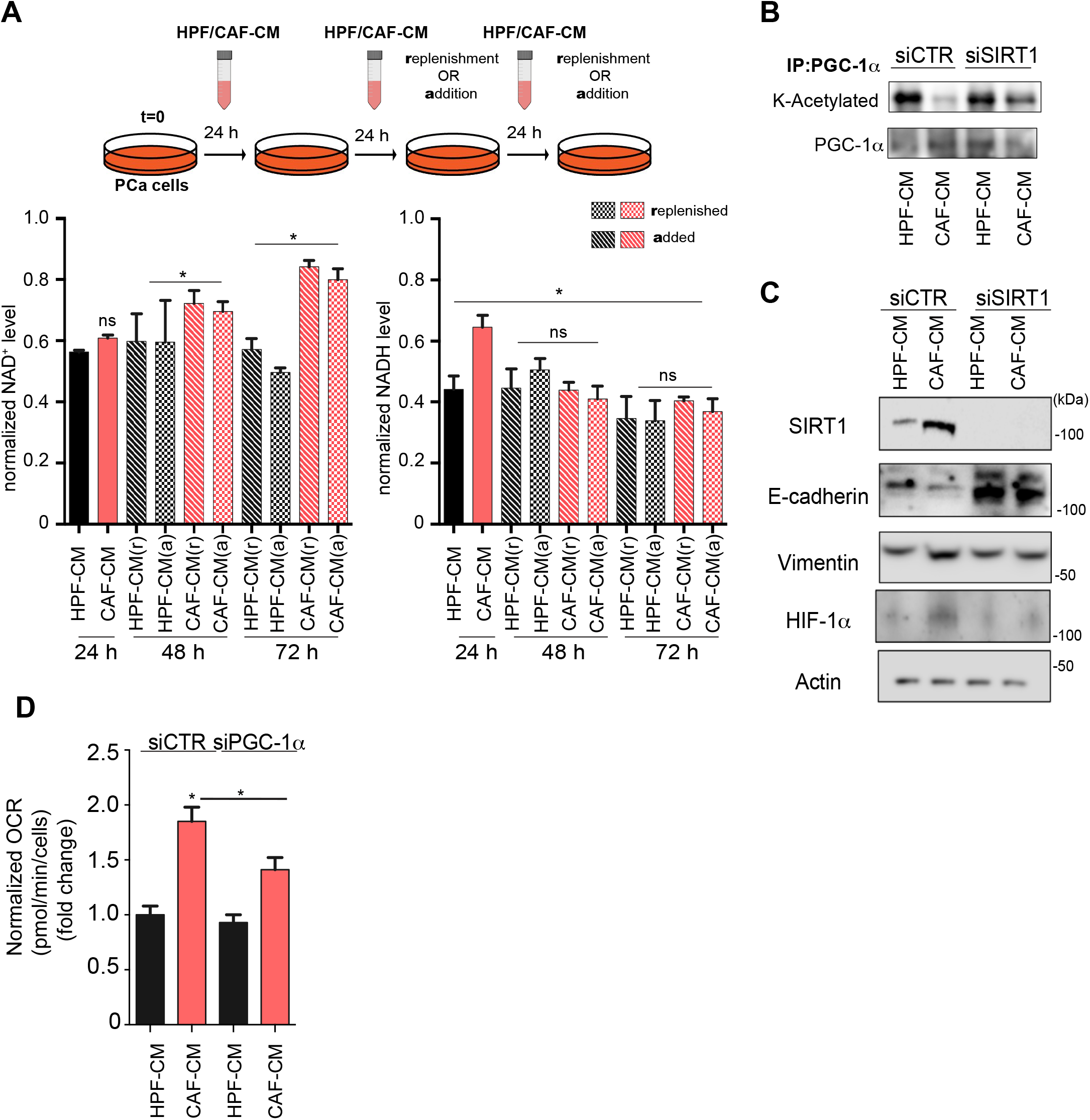
CAF-induced SIRT1/PGC-1α axis affects EMT and mitochondrial metabolism in PCa cells. (A) Normalized NAD^+^ and NADH levels were assessed in PC3 cells, by replenishing (r) or adding (a) HPF-CM or CAF-CM every 24 h, as depicted in the experimental setting above. Mean ± SEM in (C), *n* = 3. ns, no significant, **P < 0.05, by one-way ANOVA, with Tukey post hoc tests. (B) Immunoprecipitation and acetylation status of PGC-1α were assessed in PC3 silenced for SIRT1 and treated with HPF-or CAF-CM, as indicated. (C) Total lysates from DU145 cells silenced for SIRT1 were subjected for WB analysis. (D) Measurement of OCR in PC3 cells silenced for PGC-1α and treated with HPF-or CAF-CM.

**Figure S2.**
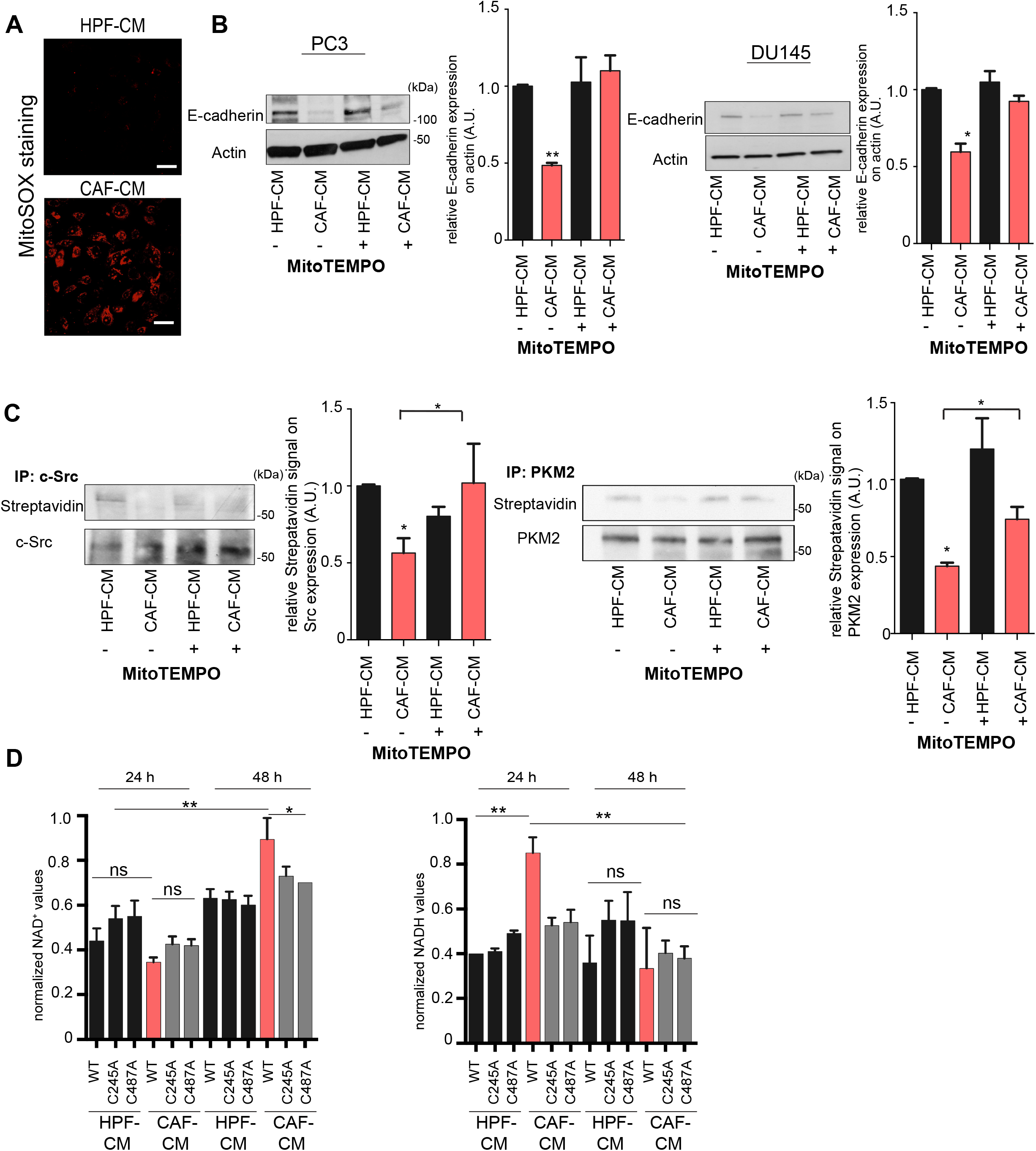
CAFs-induced mtROS generation drives Src/PKM2 oxidation and altering such redox status impairs CAF-associated EMT engagement in prostate cancer cells. (A) Representative images of mtROS levels in PC3 cells exposed to HPF-or CAF-CM. mtROS were evaluated by MitoSOX staining and analyzed by confocal fluorescence analysis. Scale bar, 10 μm. B-C. E-cadherin expression and Src/PKM2 oxidation were assessed in prostate cancer cells by WB analysis on total cell lysates or Src/PKM2 immunoprecipitations from BIAM-treated lysates, respectively. MitoTEMPO was used as mtROS scavenger. Quantification plots of E-cadherin and streptavidin signals normalized on actin, c-Src or PKM2 from three independent experiments were shown. (D) Normalized NAD^+^ and NADH levels were assessed in PC3 cells, transfected with redox-insensitive Src mutants, and then exposed to HPF-or CAF-CM for 24 h and 48 h. Data represent mean ± SEM, *n* = 3. **P < 0.05; **P < 0.005, by one-way ANOVA, with Tukey post hoc tests.

**Figure S3.**
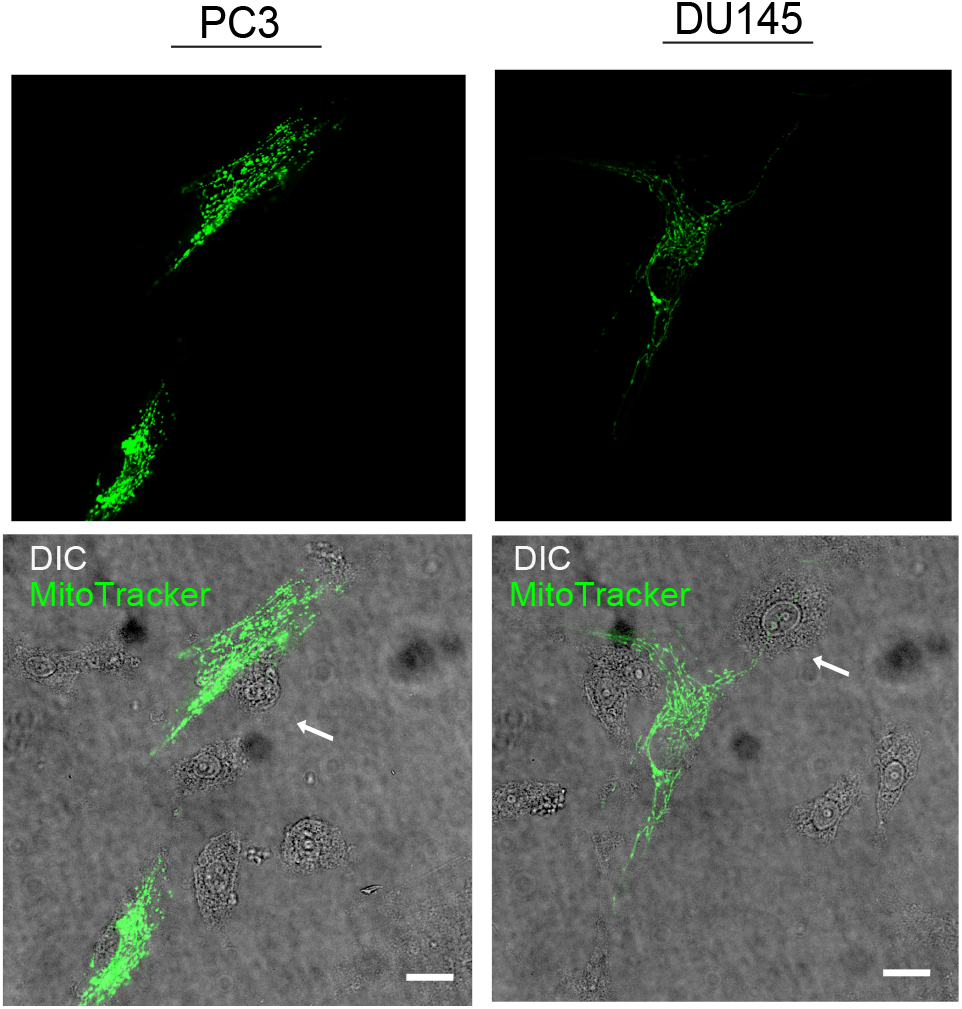
HPF cocultered with PC3 and DU145 cells do not supply own mitochondria. Representative images of HPFs labeled with MitoTracker Green cocultered with unstained PCa cells, indicated with white arrows. Scale bar, 10 μm

## REFERENCES

1. Brauer HA, Makowski L, Hoadley KA, Casbas-Hernandez P, Lang LJ, Romàn-Pèrez E et al. Impact of tumor microenvironment and epithelial phenotypes on metabolism in breast cancer. Clin Cancer Res 2013; 19: 571–585.

2. Fiaschi T, Marini A, Giannoni E, Taddei ML, Gandellini P, De Donatis A et al. Reciprocal metabolic reprogramming through lactate shuttle coordinately influences tumor-stroma interplay. Cancer Res 2012; 72: 5130–5140.

3. Giannoni E, Taddei ML, Morandi A, Comito G, Calvani M, Bianchini F et al. Targeting stromal-induced pyruvate kinase M2 nuclear translocation impairs oxphos and prostate cancer metastatic spread. Oncotarget 2015; 6: 24061–24074.

4. Ippolito L, Marini A, Cavallini L, Morandi A, Pietrovito L, Pintus G et al. Metabolic shift toward oxidative phosphorylation in docetaxel resistant prostate cancer cells. Oncotarget 2016.

5. Chandel NS. Evolution of Mitochondria as Signaling Organelles. Cell Metab 2015; 22: 204–206.

6. Viale A, Pettazzoni P, Lyssiotis CA, Ying H, Sánchez N, Marchesini M et al. Oncogene ablation-resistant pancreatic cancer cells depend on mitochondrial function. Nature 2014; 514: 628–632.

7. Weinberg F, Hamanaka R, Wheaton WW, Weinberg S, Joseph J, Lopez M et al. Mitochondrial metabolism and ROS generation are essential for Kras-mediated tumorigenicity. Proc Natl Acad Sci U S A 2010; 107: 8788–8793.

8. LeBleu VS, O’Connell JT, Gonzalez Herrera KN, Wikman H, Pantel K, Haigis MC et al. PGC-1α mediates mitochondrial biogenesis and oxidative phosphorylation in cancer cells to promote metastasis. Nat Cell Biol 2014; 16: 992–1003, 1001–1015.

9. Torrano V, Valcarcel-Jimenez L, Cortazar AR, Liu X, Urosevic J, Castillo-Martin M et al. The metabolic co-regulator PGC1α suppresses prostate cancer metastasis. Nat Cell Biol 2016; 18: 645–656.

10. Tan AS, Baty JW, Dong LF, Bezawork-Geleta A, Endaya B, Goodwin J et al. Mitochondrial genome acquisition restores respiratory function and tumorigenic potential of cancer cells without mitochondrial DNA. Cell Metab 2015; 21: 81–94.

11. Dong LF, Kovarova J, Bajzikova M, Bezawork-Geleta A, Svec D, Endaya B et al. Horizontal transfer of whole mitochondria restores tumorigenic potential in mitochondrial DNA-deficient cancer cells. Elife 2017; 6.

12. Spees JL, Olson SD, Whitney MJ, Prockop DJ. Mitochondrial transfer between cells can rescue aerobic respiration. Proc Natl Acad Sci U S A 2006; 103: 1283–1288.

13. Ishikawa K, Takenaga K, Akimoto M, Koshikawa N, Yamaguchi A, Imanishi H et al. ROS-generating mitochondrial DNA mutations can regulate tumor cell metastasis. Science 2008; 320: 661–664.

14. Guarente L. Calorie restriction and sirtuins revisited. Genes Dev 2013; 27: 2072–2085.

15. Houtkooper RH, Pirinen E, Auwerx J. Sirtuins as regulators of metabolism and healthspan. Nat Rev Mol Cell Biol 2012; 13: 225–238.

16. Gambini J, Gomez-Cabrera MC, Borras C, Valles SL, Lopez-Grueso R, Martinez-Bello VE et al. Free [NADH]/[NAD(+)] regulates sirtuin expression. Arch Biochem Biophys 2011; 512: 24–29.

17. Rodgers JT, Lerin C, Haas W, Gygi SP, Spiegelman BM, Puigserver P. Nutrient control of glucose homeostasis through a complex of PGC-1alpha and SIRT1. Nature 2005; 434: 113–118.

18. Nemoto S, Fergusson MM, Finkel T. SIRT1 functionally interacts with the metabolic regulator and transcriptional coactivator PGC-1{alpha}. J Biol Chem 2005; 280: 16456–16460.

19. Giannoni E, Bianchini F, Calorini L, Chiarugi P. Cancer associated fibroblasts exploit reactive oxygen species through a proinflammatory signature leading to epithelial mesenchymal transition and stemness. Antioxid Redox Signal 2011; 14: 2361–2371.

20. Selak MA, Armour SM, MacKenzie ED, Boulahbel H, Watson DG, Mansfield KD et al. Succinate links TCA cycle dysfunction to oncogenesis by inhibiting HIF-alpha prolyl hydroxylase. Cancer Cell 2005; 7: 77–85.

21. MacKenzie ED, Selak MA, Tennant DA, Payne LJ, Crosby S, Frederiksen CM et al. Cell-permeating alpha-ketoglutarate derivatives alleviate pseudohypoxia in succinate dehydrogenase-deficient cells. Mol Cell Biol 2007; 27: 3282–3289.

22. Weinberg F, Chandel NS. Mitochondrial metabolism and cancer. Ann N Y Acad Sci 2009; 1177: 66–73.

23. Cannito S, Novo E, di Bonzo LV, Busletta C, Colombatto S, Parola M. Epithelial-mesenchymal transition: from molecular mechanisms, redox regulation to implications in human health and disease. Antioxid Redox Signal 2010; 12: 1383–1430.

24. Porporato PE, Payen VL, Pérez-Escuredo J, De Saedeleer CJ, Danhier P, Copetti T et al. A mitochondrial switch promotes tumor metastasis. Cell Rep 2014; 8: 754–766.

25. Dikalova AE, Bikineyeva AT, Budzyn K, Nazarewicz RR, McCann L, Lewis W et al. Therapeutic targeting of mitochondrial superoxide in hypertension. Circ Res 2010; 107: 106–116.

26. Nazarewicz RR, Dikalova A, Bikineyeva A, Ivanov S, Kirilyuk IA, Grigor’ev IA et al. Does scavenging of mitochondrial superoxide attenuate cancer prosurvival signaling pathways? Antioxid Redox Signal 2013; 19: 344–349.

27. Porporato PE, Sonveaux P. Paving the way for therapeutic prevention of tumor metastasis with agents targeting mitochondrial superoxide. Mol Cell Oncol 2015; 2: e968043.

28. Giannoni E, Buricchi F, Raugei G, Ramponi G, Chiarugi P. Intracellular reactive oxygen species activate Src tyrosine kinase during cell adhesion and anchorage-dependent cell growth. Mol Cell Biol 2005; 25: 6391–6403.

29. Rogers RS, Bhattacharya J. When cells become organelle donors. Physiology (Bethesda) 2013; 28: 414–422.

30. Rustom A, Saffrich R, Markovic I, Walther P, Gerdes HH. Nanotubular highways for intercellular organelle transport. Science 2004; 303: 1007–1010.

31. Lou E, Fujisawa S, Morozov A, Barlas A, Romin Y, Dogan Y et al. Tunneling nanotubes provide a unique conduit for intercellular transfer of cellular contents in human malignant pleural mesothelioma. PLoS One 2012; 7: e33093.

32. Pasquier J, Guerrouahen BS, Al Thawadi H, Ghiabi P, Maleki M, Abu-Kaoud N et al. Preferential transfer of mitochondria from endothelial to cancer cells through tunneling nanotubes modulates chemoresistance. J Transl Med 2013; 11: 94.

33. Caicedo A, Fritz V, Brondello JM, Ayala M, Dennemont I, Abdellaoui N et al. MitoCeption as a new tool to assess the effects of mesenchymal stem/stromal cell mitochondria on cancer cell metabolism and function. Sci Rep 2015; 5: 9073.

34. Salem AF, Whitaker-Menezes D, Lin Z, Martinez-Outschoorn UE, Tanowitz HB, Al-Zoubi MS et al. Two-compartment tumor metabolism: autophagy in the tumor microenvironment and oxidative mitochondrial metabolism (OXPHOS) in cancer cells. Cell Cycle 2012; 11: 2545–2556.

35. Morandi A, Giannoni E, Chiarugi P. Nutrient Exploitation within the Tumor-Stroma Metabolic Crosstalk. Trends Cancer 2016; 2: 736–746.

36. Chen YJ, Mahieu NG, Huang X, Singh M, Crawford PA, Johnson SL et al. Lactate metabolism is associated with mammalian mitochondria. Nat Chem Biol 2016; 12: 937–943.

37. Chen EI, Hewel J, Krueger JS, Tiraby C, Weber MR, Kralli A et al. Adaptation of energy metabolism in breast cancer brain metastases. Cancer Res 2007; 67: 1472–1486.

38. Andrzejewski S, Klimcakova E, Johnson RM, Tabariès S, Annis MG, McGuirk S et al. PGC-1α Promotes Breast Cancer Metastasis and Confers Bioenergetic Flexibility against Metabolic Drugs. Cell Metab 2017; 26: 778–787.e775.

39. Kong X, Wang R, Xue Y, Liu X, Zhang H, Chen Y et al. Sirtuin 3, a new target of PGC-1alpha, plays an important role in the suppression of ROS and mitochondrial biogenesis. PLoS One 2010; 5: e11707.

40. Rabinovitch RC, Samborska B, Faubert B, Ma EH, Gravel SP, Andrzejewski S et al. AMPK Maintains Cellular Metabolic Homeostasis through Regulation of Mitochondrial Reactive Oxygen Species. Cell Rep 2017; 21: 1–9.

41. Le Gal K, Ibrahim MX, Wiel C, Sayin VI, Akula MK, Karlsson C et al. Antioxidants can increase melanoma metastasis in mice. Sci Transl Med 2015; 7: 308re308.

42. Faubert B, Li KY, Cai L, Hensley CT, Kim J, Zacharias LG et al. Lactate Metabolism in Human Lung Tumors. Cell 2017; 171: 358–371.e359.

43. Aspuria PJ, Lunt SY, Väremo L, Vergnes L, Gozo M, Beach JA et al. Succinate dehydrogenase inhibition leads to epithelial-mesenchymal transition and reprogrammed carbon metabolism. Cancer Metab 2014; 2: 21.

44. Sciacovelli M, Gonçalves E, Johnson TI, Zecchini VR, da Costa AS, Gaude E et al. Fumarate is an epigenetic modifier that elicits epithelial-to-mesenchymal transition. Nature 2016; 537: 544–547.

45. Martinez-Outschoorn UE, Trimmer C, Lin Z, Whitaker-Menezes D, Chiavarina B, Zhou J et al. Autophagy in cancer associated fibroblasts promotes tumor cell survival: Role of hypoxia, HIF1 induction and NFκB activation in the tumor stromal microenvironment. Cell Cycle 2010; 9: 3515–3533.

46. Pavlides S, Whitaker-Menezes D, Castello-Cros R, Flomenberg N, Witkiewicz AK, Frank PG et al. The reverse Warburg effect: aerobic glycolysis in cancer associated fibroblasts and the tumor stroma. Cell Cycle 2009; 8: 3984–4001.

47. Giannoni E, Bianchini F, Masieri L, Serni S, Torre E, Calorini L et al. Reciprocal activation of prostate cancer cells and cancer-associated fibroblasts stimulates epithelial-mesenchymal transition and cancer stemness. Cancer Res 2010; 70: 6945–6956.

48. Santi A, Caselli A, Ranaldi F, Paoli P, Mugnaioni C, Michelucci E et al. Cancer associated fibroblasts transfer lipids and proteins to cancer cells through cargo vesicles supporting tumor growth. Biochim Biophys Acta 2015; 1853: 3211–3223.

49. Benton CR, Yoshida Y, Lally J, Han XX, Hatta H, Bonen A. PGC-1alpha increases skeletal muscle lactate uptake by increasing the expression of MCT1 but not MCT2 or MCT4. Physiol Genomics 2008; 35: 45–54.

50. Summermatter S, Santos G, Pérez-Schindler J, Handschin C. Skeletal muscle PGC-1α controls whole-body lactate homeostasis through estrogen-related receptor α-dependent activation of LDH B and repression of LDH A. Proc Natl Acad Sci U S A 2013; 110: 8738–8743.

51. Pittelli M, Felici R, Pitozzi V, Giovannelli L, Bigagli E, Cialdai F et al. Pharmacological effects of exogenous NAD on mitochondrial bioenergetics, DNA repair, and apoptosis. Mol Pharmacol 2011; 80: 1136–1146.

52. Frezza C, Cipolat S, Scorrano L. Organelle isolation: functional mitochondria from mouse liver, muscle and cultured fibroblasts. Nat Protoc 2007; 2: 287–295.

53. Giannoni E, Fiaschi T, Ramponi G, Chiarugi P. Redox regulation of anoikis resistance of metastatic prostate cancer cells: key role for Src and EGFR-mediated pro-survival signals. Oncogene 2009; 28: 2074–2086.

